# Quantifying the Effect of Anti-cancer Compound (Piperlongumine) on Cancer Cells Using Single-Cell Force Spectroscopy

**DOI:** 10.1101/2021.09.03.458840

**Authors:** Nayara Sousa de Alcântara-Contessoto, Marinônio Lopes Cornélio, Ching-Hwa Kiang

## Abstract

Natural compounds have shown a great potential in anti-cancer research by tumor growth inhibition and anti-metastatic properties. Piperlongumine (PL) is a natural compound derived from pepper species that has been demonstrated to have anti-cancer effect on HeLa cells. Here we focus on understanding the mechanical properties of HeLa cells under PL treatment, using Atomic Force Microscopy (AFM) based single-cell manipulation technique. We used AFM to pull single HeLa cells and acquired the force-distance curves that presented stepwise patterns. We analyzed the step force (SF) and observed that cells treated with PL exhibit higher force compared to control cells. This SF increase was also observed in experiments performed on substrates of different stiffness. Therefore, analyzing SF, it is possible to investigate the effect of PL on the mechanical properties of the HeLa cells. The understanding of the PL action on HeLa cells’ mechanical properties may help in the development of effective therapeutic drugs against cancers.

## 1. Introduction

Anti-cancer research is a challenge and a worldwide interest. Cancer causes millions of people’s death every year, and in 2020, it was responsible for around 10 million [1, 2]. Cancer diseases have two main concerns: cell growth and cell migration, which can be uncontrolled [3]. Metastasis, being the cause for most cancer patients’ mortality, receives increasing attention in both scientific and clinical research [4]. Understanding cancer cell mechanics can improve the metastasis combat through observations into the diagnosis, prognosis, and treatment [4, 5]. In this sense, the mechanical properties of single molecules and single-cells have been investigated using atomic force microscopy (AFM) and optical tweezers [6, 7, 8, 9, 10, 11].

In eukaryotic cells, physical forces can act through the cytoskeleton polymers (actin filaments, microtubules, and intermediate filaments) to control the mechanical properties, adhesion forces, and cellular behavior [7, 12]. The cytoskeletal dynamic generates effects on cell motility, division, and overall mechanical processes [12, 13]. Its dysfunction may result in chromosomal instability, mitotic arrest, and cell death, becoming common and useful targets to drug design [14]. In particular, AFM can detect a range of forces from picoNewtons to nanoNewtons, and it has been a powerful tool for high-resolution imaging and mechanical measurement of single-cell investigations in near-physiological conditions [15, 13, 6]. AFM technique can also help understand how cell mechanics are affected by drugs, i.e., providing an alternative to evaluate drug-cell interactions [15, 13].

Piperlonumine (PL), also known as Piplartine (Figure 1), is a natural compound that presents several pharmacological properties, such as genotoxic, cytotoxic, antimetastatic, and antitumoral [16, 17]. Its target can be reached through the blood plasma, as observed through *in vitro* and *in vivo* studies [18, 16, 19, 20]. Recently, Meegan et al. presented PL as a microtubule-destabilizing agent with antiproliferative effects in breast cancer cells [21]. Additionally, Henrique et al. showed the inhibitory effect of PL on the α-tubulin expression in endothelial cells [22]. Microtubule-targeting agents (MTA) are potent mitotic poisons that inhibit eukaryotic cell proliferation, promote cell death by suppressing microtubule dynamic instability, and interfere with intracellular transport [5, 23, 24]. PL presents different mechanisms of action and has been proposed as a potential anti-cancer drug and an interesting compound to be investigated [16, 25, 26].

**Figure 1:**
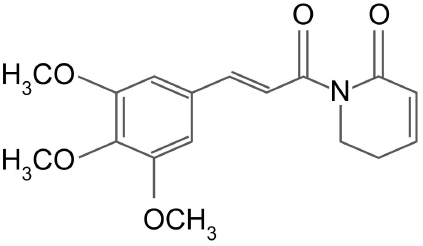
Chemical structure of Piperlongumine (PL), also known as Piplartine, (5,6-dihydro-1-[(2E) −1-oxo-3-(3,4,5-trimethoxyphenyl) −2-propen-1-yl] −2 (1H)-pyridinone) [16, 31].

Studies suggest the cytotoxicity of piperlongumine in a dose-dependence on HeLa cells [27, 25]. The literature reports the EC_50_ (concentration inhibiting cell growth by 50%) as 2.7 and 7.1 *μ*M [27, 25]. Notwithstanding the advances in understanding biochemical properties, there is a lack of information regarding the mechanical features and how they can affect cancer cell stability under the influence of drugs. AFM has been used to investigate the viscosity, surface, stiffness, adhesive properties of drug effects on cells [15, 13, 28, 29, 30]. Mechanical force is a nonspecific parameter, i.e., independent of the types of molecules, cells, and tissues [8]. Thus, it can be used to sense the influences of drugs on cells.

In this study, we performed mechanical characterization of HeLa cells in response to PL presence. The analyses were performed at different conditions of HeLa at piperlongumine treatment and their respective controls, using AFM through single-cell manipulation. We varied the PL concentrations, treatment times, and culture substrates (the surface where cells grow up) with different stiffness. This study provide quantified mechanical information of the compounds on cells, which can be related to PL’s antiproliferative effect. Therefore, investigate the cellular mechanical properties can aid cancer therapeutic and diagnostic research [4].

## 2. Materials and Methods

### 2.1. Materials and Solutions

Human cervical cancer cells (HeLa; ATCC) were cultured in a medium containing DMEM (Dulbecco’s Modification of Eagle Medium; Corning), supplemented with 10% FBS (Fetal Bovine Serum; Gibco) and 1% Pen-Strep (Penicillin/ Streptomycin; Gibco), and incubated at 37°C in a 5% CO_2_ humidified atmosphere. Piperlongumine (C_17_H_19_NO_5_) was donated by collaborators and dissolved in DMSO (Dimethylsulfoxide; ATCC) at final concentrations of 5, 10, and 15 *μ*M [16, 31]. Polymeric substrates with specific stiffness (0.5 and 16 kPa) were donated by a collaborator. MLCT-O10 cantilever (Bruker), tipless, was used as a probe in AFM experiments.

### 2.2. Cellular Sample Preparation

Sterilized steel disks were covered by glass or specific polymeric substrates (with determined stiffness) and kept in 35mm Petri dishes; the cells (10^5^ cells/ml) were subcultured with DMEM supplemented (10%FBS and 1%Pen Strep) medium on this assay and kept overnight in an incubator (5% CO_2_ and 37°C). After this step, the cells were treated by Piperlongumine at different action times and different concentrations. The equivalent volume of vehicle DMSO without PL was added to the DMSO control group and no DMSO/PL to the negative control group. DMSO at less than 0.5% in the cell culture medium [32].

### 2.3. Atomic Force Microscopy

Force-distance curves were performed using a MultiMode 8 Atomic Force Microscope (Bruker) equipped with a Nanoscope V controller (Bruker), a PicoForce (Force Spectroscopy Control Module, Digital Instruments), and an optical microscope (Digital Instruments). The cantilever (MLCT-O10) was calibrated, measuring the deflection sensitivity and using the Thermal Tune to determine the cantilever spring constant [33].

The single-cells on specific disks (glass or soft substrates), at room temperature, were analyzed using the cantilever probes with spring constant (K) approximately equal to 0.01 N/m (shape C triangular).

Figure 2 - A presents an AFM framework. The data collected by AFM contact mode was performed positioning (X-Y axis) the probe (AFM cantilever) on a single-cell. The parameters are set up (ramp size, velocities, trig threshold, delay times, spring constant) according to the experiment’s aim (Table 1). A laser beam is reflected by the AFM cantilever and collected in a photodetector (photodiode) while the probe is engaged and withdrawn from the sample (Z-axis).

**Figure 2:**
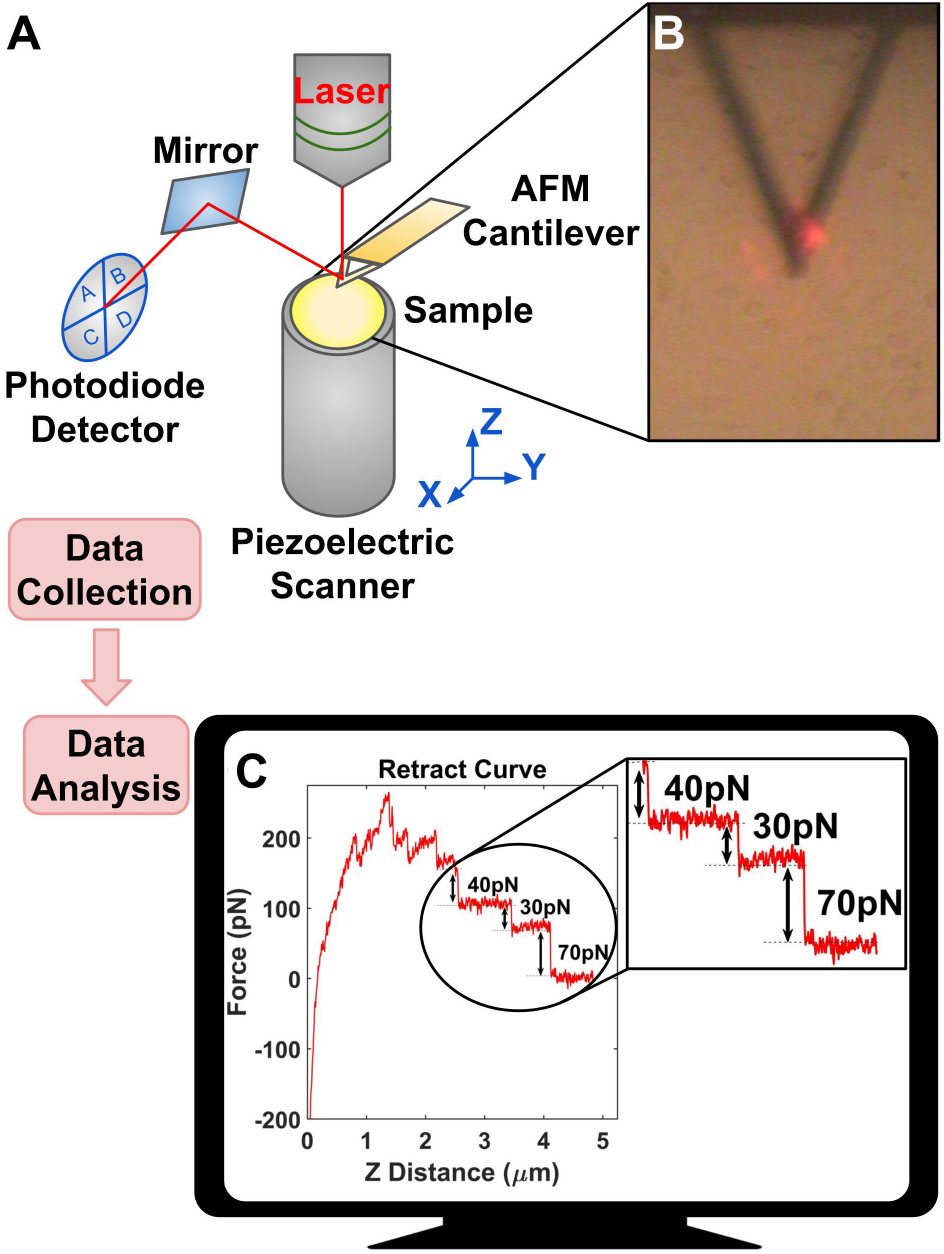
Tether Force, a schematic figure from data collection to data analysis. A - AFM framework. B - Image of the optical microscope attached to AFM: Single-cells selection in the sample. C - The output of Nanoscope software: force-distance curve from AFM cantilever retract move (half cycle) used for analyses of the tether force. The double arrows indicate the tether force (or step force) for each abrupt detaching of retracting curve. Highlighted are the specific values.

**Table 1:** Parameters for analyzes of the tether forces in contact mode AFM, using MLCT-O10 cantilever.

| Ramp<br>Size | Forward<br>Velocity | Reverse<br>Velocity | Trig<br>Threshold | Surface<br>Delay | Retracted<br>Delay | Spring<br>Constant |
| --- | --- | --- | --- | --- | --- | --- |
| 5 $\mu\text{m}$ | 3 $\mu\text{m/s}$ | 3 $\mu\text{m/s}$ | 1 nN | 3 s | 3 s | 0.01 N/m |

#### 2.3.1. Force-Distance Curves

Each cycle of the AFM probe is performed by engaging and withdrawing it from the cell surface. The cycle generates a pair of force-distance curves (approach and retract). Figure 2 - C presents a usual retracts curve.

#### 2.3.2. Tether Force Studies

Figure 2 - C presents plateaus profile occurring at constant forces, separated by steps. The multiple-step formation was observed through force-distance curves generated by an AFM when the probe withdrawal from the cell surface after its engagement. It is suggested that the pulling process results in the formation of thin nanotubes or tethers [13]. Then, similar to springs connected in parallel, multiple tethers can be formed between the cantilever and the cell [6]. The double arrows in figure 2 - C indicate the discrete step force (SF or ΔF) between consecutive plateaus, interpreted as simultaneous elongation and sequential loss of tether [13].

The tether force (or Step Force) was obtained by measuring retract curve steps (Figure 2 - C) from several single-cells. Hundreds of force-distance curves were recorded for each assay. A step-fitting algorithm was used to extract the step force values [34].

An ensemble of step force values generates distribution graphs (histogram and violin plots). The histogram is fitted by a Gaussian curve and identifies the most probable tether force (SF) value. The violin plots were generated to observe the median value and the quartiles of the data within the distribution.

The tether force experiments aim to extract biomechanical information of HeLa cells in response to PL action in different assays (action time, concentration, and culture substrate dependence). Table 1 shows the AFM setup parameters for performing the measure.

## 3. Results and Discussions

### 3.1. Validation Method

The investigated conditions were monitored to guarantee that the signal’s origin is only due to the influence of the compound. Figure 3 - A presents step force analyzes for cells with DMSO at different times. DMSO presence did not affect the tether force for HeLa cells, as observed during 30 hours of the administration. The most probable SF value was kept at around 50pN (Figure 3 - A). Therefore, it is assumed that the medium of compound solubilization (DMSO) does not interfere in its action.

**Figure 3:**
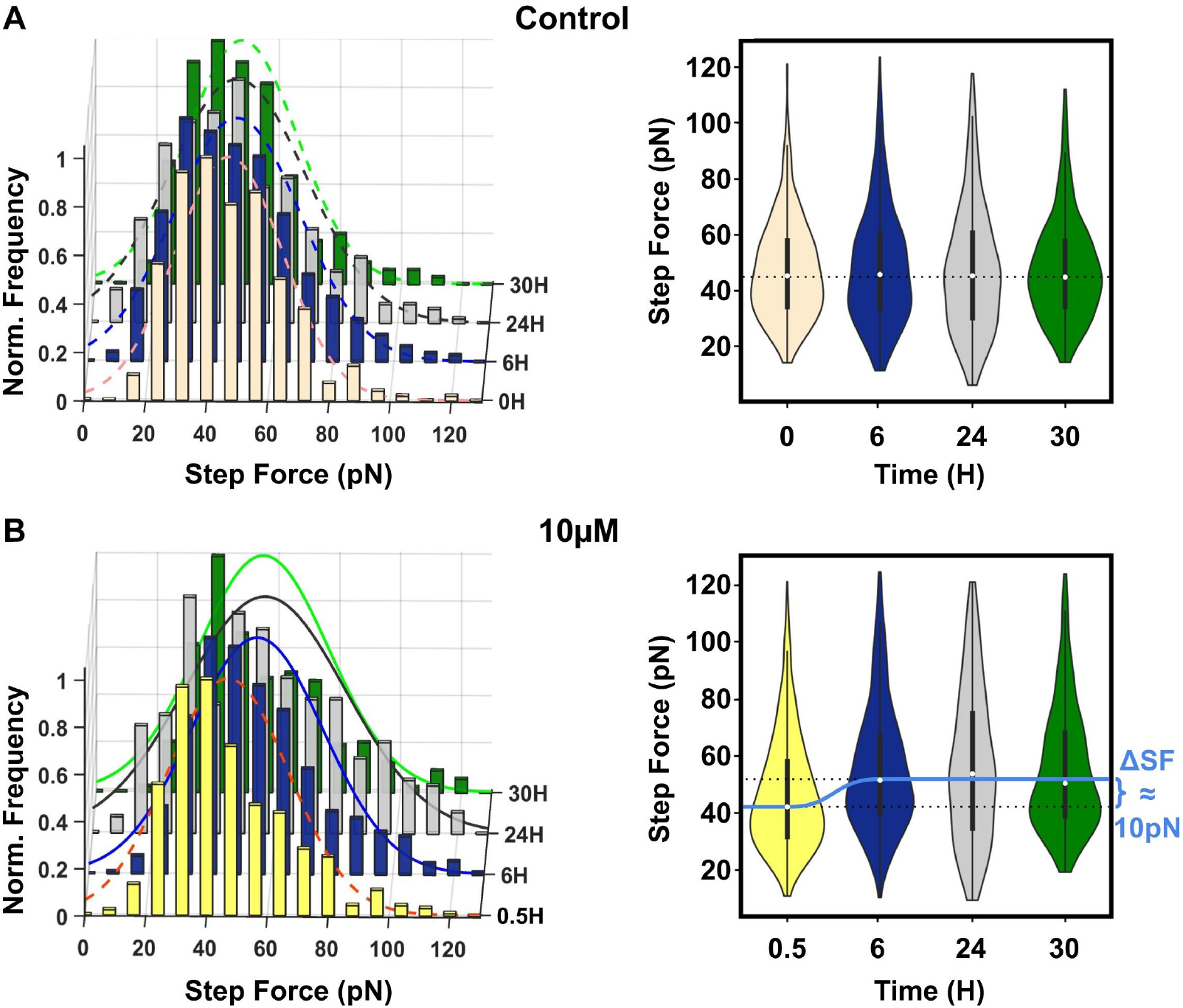
Histograms (left) and violin plots (right) of step force from retract force-distance curves of HeLa cells at time-dependence. A - Control cells (without (0 h) and with DMSO (6, 24, and 30 hours) at 0*μ*M of PL). The mean of the histograms is around 50pN (dashed curves). B - Treated cells with PL (10*μ*M) at 0.5, 6, 24, and 30 hours. The mean of the histograms is around 50pN (dashed curve) and 60pN (continuous curves) for the 0.5 hour and the other analyzed times (6, 24, and 30 hours), respectively. The black bar (right figures) presents the first and third quartile of the data. The white dots (right figures) present the median value for SF. ΔSF is the variation of the step force’s median value after 6 hours of treatment with PL (10*μ*M) until the 30 hours observed (the light blue curve is to guide the eyes (Figure 3-B, right)). The cells were cultured on glass substrates. The statistical analyzes are presented in Tables S1 and S2 of Supporting Information.

Furthermore, Figure 3 - B presents step force analyzes for the compound action time. The most probable SF value for HeLa at the first 30 minutes of piperlongumine action (10*μ*M, SF ≈ 50pN) is equal to the control group analyses (Figure 3 - A). After 6 hours of piperlongumine treatment (10*μ*M, SF ≈ 60pN), the distribution peak shifted to greater step force than the control group (ΔSF ≈ 10pN).

Bezerra et al. presented several PL’s mechanisms of action, such as the increase of reactive oxygen species (ROS) [16, 25]. Recently, it was observed a correlation between ROS-free environment and dynamic instability, which has been related as important in cell division and motility [35, 5, 23]. Micro tubules showed an enhanced dynamic instability in a ROS-free environment [35]. PL has been reported to increase the level of ROS in HeLa cells within 1.5 hours [25]. The increase of ROS level by PL on HeLa cells appears to be related to suppressing dynamic instability on microtubules, and it can be associated with the step force variation (ΔSF) by PL action time on HeLa (Figure 3 - B). Figure 4 suggests that the mechanical effect of piperlongumine on HeLa cells occurs at the first 6 hours of treatment. The time-dependence results indicate that tether force analyzes on cells are sensitive to drug action time.

**Figure 4:**
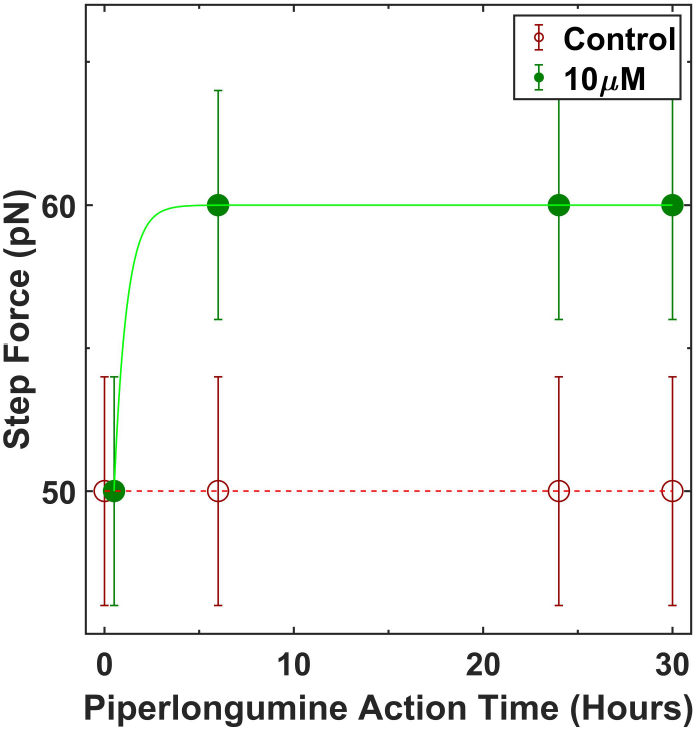
Time-dependent, most probable step force. HeLa cells in the presence of 10*μ*M of PL (30 min, 6, 24, and 30 Hours) and control (0, 6, 24, and 30 Hours). The red and green curves are to guide the eyes.

### 3.2. Concentration Influences

Piperlongumine is reported to be cytotoxic in a concentration dependence [27, 25]. In this sense, we analyzed the biomechanical effects dependents of the piperlongumine concentration on HeLa cells.

Figure 5 presents the step force distributions for HeLa cells treated with 5, 10, and 15*μ*M of piperlongumine, compared with the control (DMSO), for a constant action time (24 hours). The cells were not sensitive to step force changes in 5*μ*M of piperlongumine as compared to control, i.e., no shift at most probable SF was observed for this concentration. However, in treatment with 10*μ*M and 15*μ*M, the distributions showed a specific increase of step force value, for each concentration, compared to the control (ΔSF ≈ 10pN).

**Figure 5:**
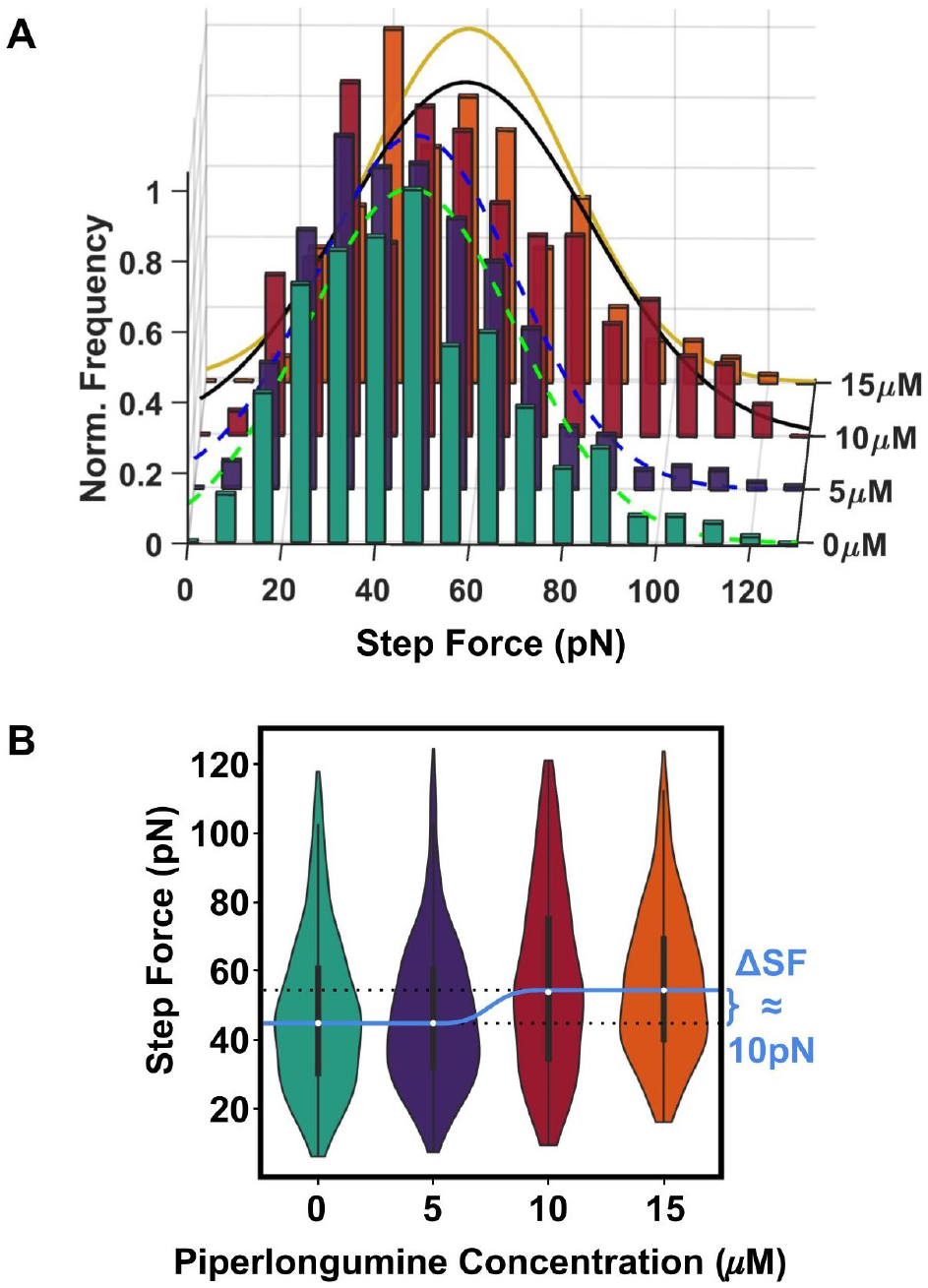
A - Histograms of step force from retract force-distance curves and B - Violin plots of step force of HeLa cells at concentration-dependence (0 (control), 5, 10, and 15*μ*M) and 24 hours of treatment. The median values (white dots) increase around 10pN (ΔSF, Figure 5-B) between 5*μ*M and 10*μ*M. The light blue curve is to guide the eyes, and the black bar presents the first and third quartile of the data (Figure 5-B). The cells were cultured on glass substrates. The statistical analysis is presented in Table S3 of Supporting Information.

The biomechanical effects of PL on HeLa cells is sensitive by the tether force analyzes in a concentration-dependence. These results can be related to the cellular viability value in 50% for HeLa cells in PL presence, which are presented in the literature as smaller than 8*μ*M [36, 27, 25].

### 3.3. Extracellular Environment Influences

Studies reported that the microenvironment of the cell culture influences the mode and dynamics of cancer cell invasion [3, 6]. Here, it was investigated the step force of HeLa cells in different substrates of cellular culture with specific stiffness (0.5kPa, 16kPa, and glass). Figure 6 presents a shift of distribution peak to larger step force value, from control to piperlongumine treated cells, for each specific substrate.

**Figure 6:**
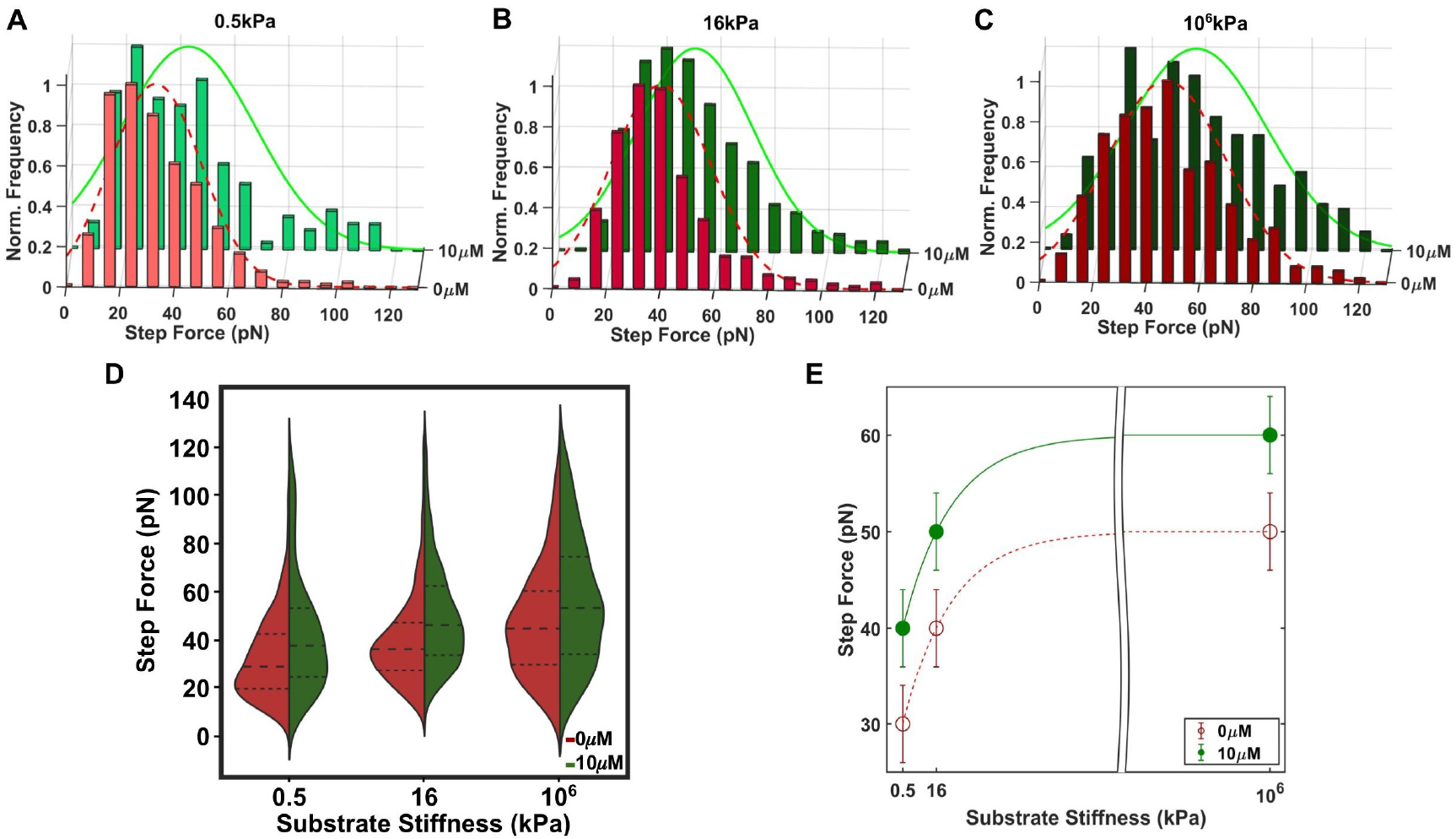
A to C - Histograms of step force from retract force-distance curves and D - Violin plots of the step force for HeLa cells at matrix-dependence (0.5kPa, 16kPa, and 10^6^kPa (glass)) in the presence of PL (10 *μ*M) and control (0 *μ*M of PL) after 24 hours of treatment. The median values (black dashed lines) increase as the matrix (substrate) is stiffer. Besides, the median values increase between control (red half-violin) and 10 *μ*M (green half-violin) in each specific matrix substrate. The black dotted lines present the first and third quartiles of the data. E - The most probable step force (round value) of HeLa cells at matrix-dependence (0.5kPa, 16kPa, and glass) in the presence of PL (10 *μ*M), green curve, and control (dashed red curve) after 24 hours of treatment. The statistical analysis is presented in Table S4 of Supporting Information.

Figure 6 - E shows the most probable step force for HeLa cells in different substrates matrices. The SF values at the same condition, either control or 10*μ*M of piperlongumine, increase with the stiffer substrate (as guided by the lines in figure 6 - E). These results indicate the extracellular environment influences the biomechanical effects.

Furthermore, for each substrate analyzed, the variation of the most probable SF value between control and treated cells is the same, and around 10pN (ΔSF_10*μM–C*_, Table S5 of Supporting Information). These analyses indicate that the effect of PL on HeLa cells presents a ΔSF pattern that is independent of the substrate rigidity.

The curve in figure 6 - E (Table S6 of Supporting Information) of SF for different substrates (0.5kPa, 16kPa, and glass) was analyzed using Equation 1 for each sample (Control and treated with 10*μ*M of PL).

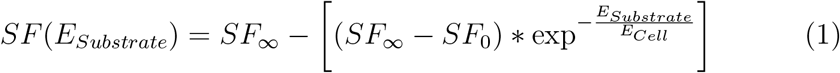

SF^∞^ and SF_0_ are the extrapolated step force values (SF) to infinity and zero, respectively; E*_Substrate_* is the substrate stiffness; E*_Cell_* is the cell stiffness [6].

As presented in the literature, the tether properties depend on actin filaments and microtubules network, which are the major cytoskeleton components [9, 13]. Recently, Pontes et al. observed the presence of the actin filaments in tether structure, besides the membrane [10, 11]. In previous studies, cells treated with actin microfilaments-destabilizing agent (latrunculin A (LATA)), as well as glycocalyx backbone-disrupt agent (hyaluronidase), trend a decrease of the step force value in the presence of each drug [13]. However, in this study, HeLa cells treated with PL, a microtubule-destabilizing agent, presented an increase of the steps (tether) force in several assays (action times, drug concentrations, and culture substrates). Due to this different behavior, we suggest that PL should act in the non-peripheral cellular region differently from LATA and hyaluronidase. Although both PL and LATA present cytoskeleton-destabilizing properties, they target different cytoskeleton subunits (tubulin and actin, respectively) [13, 21, 37, 38, 5]. Therefore, this study suggests that tether (step) force can be used as a mechanical biomarker sensitive to the site of the cellular target by drugs.

## 4. Conclusion

Understanding the biomechanical properties in cells upon compound interaction can help elucidate the underlying mechanism of anti-cancer drug activities. In this context, Atomic Force Microscopy (AFM) is a technique that can aid the studies of drug action mechanisms quantitatively. Here, we employed AFM experiments to investigate cancer cells’ biomechanical properties under the influences of an anti-cancer compound. Different conditions of the compound and culture substrate rigidity were explored using HeLa cells. The results indicated that the step force (SF) is sensitive to the drug action time; piperlongumine acts on HeLa cells in the first 6 hours of the treatment. Additionally, SF is sensitive to the compound concentration; between 5 and 10 *μ*M of piperlongumine treatment, HeLa cells experiments present an increase of SF, a variation of around 10 pN. Besides action time and the compound concentration, SF is also sensitive to the cytoskeleton changes; HeLa cells in the presence of 10 *μ*M of the piperlongumine increase the tether force (ΔSF_10*μ M–c*_ ≈ 10pN) compared with the control, independently of the substrate stiffness. Recently, piperlongumine has been described in the literature as a microtubule-destabilizing agent [21]. In this study, we observed an increase in the step force values of Hela cell experiments in the presence of the piperlongumine in different assays (action times, drug concentrations, and culture substrates). Such SF increment suggests that piperlongumine acts by targeting the microtubule of HeLa cells. The pipeline of AFM experiments presented here showed an effective procedure to characterize the overall interactions between anti-cancer drugs and cancer cell lines.

## 5. Acknowledgments

The authors thank Ian Lian for the polymeric substrates provided, Daniel Pereira Bezerra and José Maria Barbosa-Filho for piperlongumine provided, Jingqiang Li for laboratory technical support and Lucas Fugikawa Santos for the valuable discussions. N.S.A.C. thank Conselho Nacional de Desen-volvimento Cientéfico e Tecnolégico (CNPq), Brazil, (Grant 141714/2017-4 and 163893/2018-7). C.-H.K. and N.S.A.C. thank Welch Foundation, United States, (C-1632).

